# CrOssing fiber Modeling in the Peritumoral Area using dMRI (COMPARI)

**DOI:** 10.1101/2023.05.07.539770

**Authors:** Ehsan Golkar, Drew Parker, Steven Brem, Fabien Almairac, Ragini Verma

## Abstract

Visualization of fiber tracts around the tumor is critical for neurosurgical planning and preservation of crucial structural connectivity during tumor resection. Biophysical modeling approaches estimate fiber tract orientations from differential water diffusivity information of diffusion MRI. However, the presence of edema and tumor infiltration presents a challenge to visualize crossing fiber tracts in the peritumoral region. Previous approaches proposed free water modeling to compensate for the effect of water diffusivity in edema, but those methods were limited in estimating complex crossing fiber tracts. We propose a new cascaded multi-compartment model to estimate tissue microstructure in the presence of edema and pathological contaminants in the area surrounding brain tumors. In our model (COMPARI), the isotropic components of diffusion signal, including free water and hindered water, were eliminated, and the fiber orientation distribution (FOD) of the remaining signal was estimated. In simulated data, COMPARI accurately recovered fiber orientations in the presence of extracellular water. In a dataset of 23 patients with highly edematous brain tumors, the amplitudes of FOD and anisotropic index distribution within the peritumoral region were higher with COMPARI than with a recently proposed multi-compartment constrained deconvolution model. In a selected patient with metastatic brain tumor, we demonstrated COMPARI’s ability to effectively model and eliminate water from the peritumoral region. The white matter bundles reconstructed with our model were qualitatively improved compared to those of other models, and allowed the identification of crossing fibers. In conclusion, the removal of isotropic components as proposed with COMPARI improved the bio-physical modeling of dMRI in edema, thus providing information on crossing fibers, thereby enabling improved tractography in a highly edematous brain tumor. This model may improve surgical planning tools to help achieve maximal safe resection of brain tumors.

## Introduction

Diffusion MRI (dMRI) is a non-invasive imaging technique that measures the diffusion of water molecules in the brain’s white matter. For almost three decades, dMRI tractography has improved neuroscience by providing comprehensive white matter (WM) tractography of the human brain. In clinical practice, it could aid in neurosurgical planning by enabling the visualization of fiber tracts around the tumor [1–3]. Indeed, by providing crucial information on the position and shape of white matter (WM) tracts surrounding brain tumors, this tool could be part of a therapeutic arsenal that aims at extending the tumor resection [4]while preserving structural connectivity, and hence the associated function [5]. However, the anatomical accuracy and reliability of fiber tracking is a major concern in neurosurgical applications and is highly dependent on the acquisition and processing of dMRI data [6].

The diffusion signal obtained from dMRI is fitted to biophysical models that describe the underlying tissue microstructure, allowing for the estimation of fiber orientation and connectivity. The directionality and magnitude of water diffusion are influenced by the microstructure of the brain tissue, such as the presence of cell membranes and myelin sheaths. In the case of tumors, water diffusivity is affected by the presence of edema and infiltration in the peritumoral region, which especially challenging in complex WM regions with crossing fibers leading to inaccuracies in the estimation of WM tracts. Thus, there is a need to model tissue microstructure in the presence of pathological contaminants and complex microstructure, so that the subsequent tractography reflects the underlying anatomy, facilitating surgical decisions. Thus, the overarching goal of this paper is to design a multi-compartment model that will enable tracking through crossing fiber regions in the presence of peritumoral edema.

In our previous work, FERNET [7], we modeled the peritumoral edema using a bi-tensor model with free-water and anisotropic compartments, fitted to single-shell data. Although FERNET enhanced tractography in the peritumoral area, as a single-tensor model, it was unable to model crossing fibers. There are dMRI models for tractography that fit a fixed number of fiber populations per voxel [8, 9] which have not been tested in the peritumoral region. Recently, dMRI models that facilitate tracking through crossing regions and allow for any number of fibers in each voxel using constrained spherical deconvolution (CSD) have been developed, but these also have not been tested in the peritumoral region. In these methods, including multi-shell multi-tissue CSD (MSMT-CSD) [10]], which models “CSF-like” and “GM-like” signals in addition to the WM fiber orientation distribution (FOD), the response function of WM is estimated from the highest fractional anisotropy (FA) voxels in the brain, usually in corpus callosum, and is fixed for the entire image. The assumption that one response function can be applied to all WM, however, may not hold in pathological WM tissue. In [11, 12], special dMRI acquisitions were designed to enable modeling of WM microstructure with unprecedented detail, but their model did not estimate fiber orientations and the acquisition was not clinically feasible, and therefore the method cannot be used for tractography.

Consequently, to address these shortcomings in current literature, and in acknowledgement of the need to model the peritumoral tissue for better tractography, we have developed a new tissue modeling approach for clinically feasible product acquisitions. COMPARI (CrOssing fiber Modeling in the Peritumoral Area using clinically feasible dMRI) provides an improved estimation of crossing fibers in the peritumoral region, facilitating better identification of crossing fibers and hence improved tractography. This has been achieved by employing multiple isotropic components, to better isolate the underlying anisotropic microstructure of peritumoral tissue. We first evaluate our proposed method on simulated data. We then demonstrate enhanced modeling of crossing fibers in the peritumoral area and improved tractography in brain tumor patients.

## Methods

### Multi-compartment model (COMPARI – CrOssing fiber Modeling in the Peritumoral Area using clinically feasible dMRI)

We propose a cascaded model, comprising an initial isotropic fitting, followed by multi-compartment modeling of the tissue into isotropic and anisotropic compartments. With the fractions of the compartments fixed, we finally estimate an FOD with multi-compartment spherical harmonics fitting [13].

We fit a multi-compartment (MC) consisting of two bundle compartments: an isotropic bundle and an anisotropic bundle. The isotropic bundle consists of two ball compartments corresponding to free water and hindered water. The anisotropic bundle consists of stick and zeppelin compartments, as described in [14], corresponding to restricted intra-axonal and hindered extra-axonal diffusivities, respectively.

Mathematically, the four compartments are modeled by:

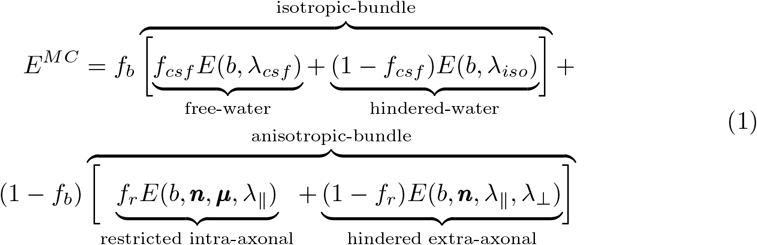

where b is the b-value,*E*(*b, λ*_*csf*_) = exp(−*bλ*_*csf*_) denotes the isotropic Gaussian signal attenuation of free water with *λ*_*csf*_ = 3 × 10^−9^*m*^2^*/s*, and *E*(*b, λ*_*iso*_) = exp(−*bλ*_*iso*_), represents isotropic signal attenuation of hindered water with magnitude *λ*_*iso*_. *λ*_*iso*_ is estimated by fitting a model with a single isotropic ball with diffusivity between 0.1× 10^−9^*m*^2^*/s* and 3× 10^−9^*m*^2^*/s*. The free parameters *f*_*b*_, *f*_*csf*_ and (1 −*f*_*csf*_) are the fractions of the overall isotropic bundle, free water and hindered fractions, respectively. The anisotropic bundle, whose volume fraction is represented by (1 − *f*_*b*_), contains a stick and a zeppelin with *f*_*r*_ and (1 − *f*_*r*_) representing the fractions of restricted intra-axonal and hindered extra-axonal diffusion, respectively.

The stick is an anisotropic cylinder model with negligible diameter represented by *E*(*b, n, µ, λ*_∥_) = exp(−*bλ*_∥_(***n***^*T*^ ***µ***)^2^), where *λ*_∥_ is the magnitude of Gaussian diffusion along ***µ*** orientation, and ***n*** represents the diffusion gradient vector. In our model, the *λ*_∥_ is fixed and equal in both stick and zeppelin, and is estimated by the voxel selection method described in [15], which automatically chooses high FA voxels, forming a reference for WM. In contrast, the hindered extra-axonal diffusion is modeled as a symmetric Gaussian distribution (zeppelin) represented by

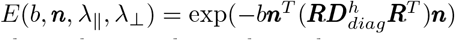 where ***R*** denotes the rotation matrix which aligns the zeppelin to the stick orientation *µ, λ*_∥_ > = *λ*_⊥_, and 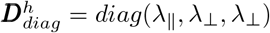.

We impose a tortuosity constraint between the diffusivity values of the zeppelin and *f*_*r*_, the fraction of the restricted intra-axonal (stick) portion of the bundle, with *λ*_⊥_ = (1 −*f*_*r*_)*λ*_∥_ [16]. Finally, we apply a condition whereby if *λ*_*iso*_ < = *λ*_∥_, the fraction *f*_*csf*_ is set to 1 and the isotropic compartment simplifies to one ball representing free water.

Next, we estimate coefficients ***c*** of a spherical harmonic function 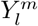(*θ, ϕ* of degree *l* and order *m*, where *θ* ∈ [0, *π*] is the inclination angle and *ϕ* ∈ [0, 2*π*] is the azimuthal angle in polar coordinates, utilizing CSD as in the multi-compartment spherical harmonics models (MC-SH) described in [10]. The Fiber Orientation Distribution (FOD) is represented by the truncated spherical harmonic series

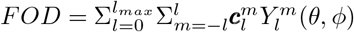 of maximum harmonic order *l*_*max*_. The final model is expressed as

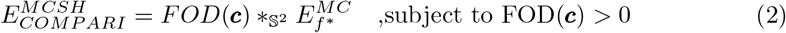

where *f* *indicates that the fitted parameters of Equation 1 are fixed, and *_𝕊_^2^ describes convolution on the sphere. The number of acquired gradient directions defines the maximum harmonic order *l*_*max*_. However, estimating higher *l*_*max*_ is possible by knowing prior information of required spherical harmonic coefficients of *l*_*max*_, assuming the non-existing directions to be zero. This is the super-CSD approach as defined in [17], and we use the regularization parameter (*λ* = 1) and threshold orientation density (*τ* = ten percent of the mean initial FOD amplitude) which were shown to be optimal [18]. The schema of our proposed method is shown in Fig. 1

**Fig 1.**
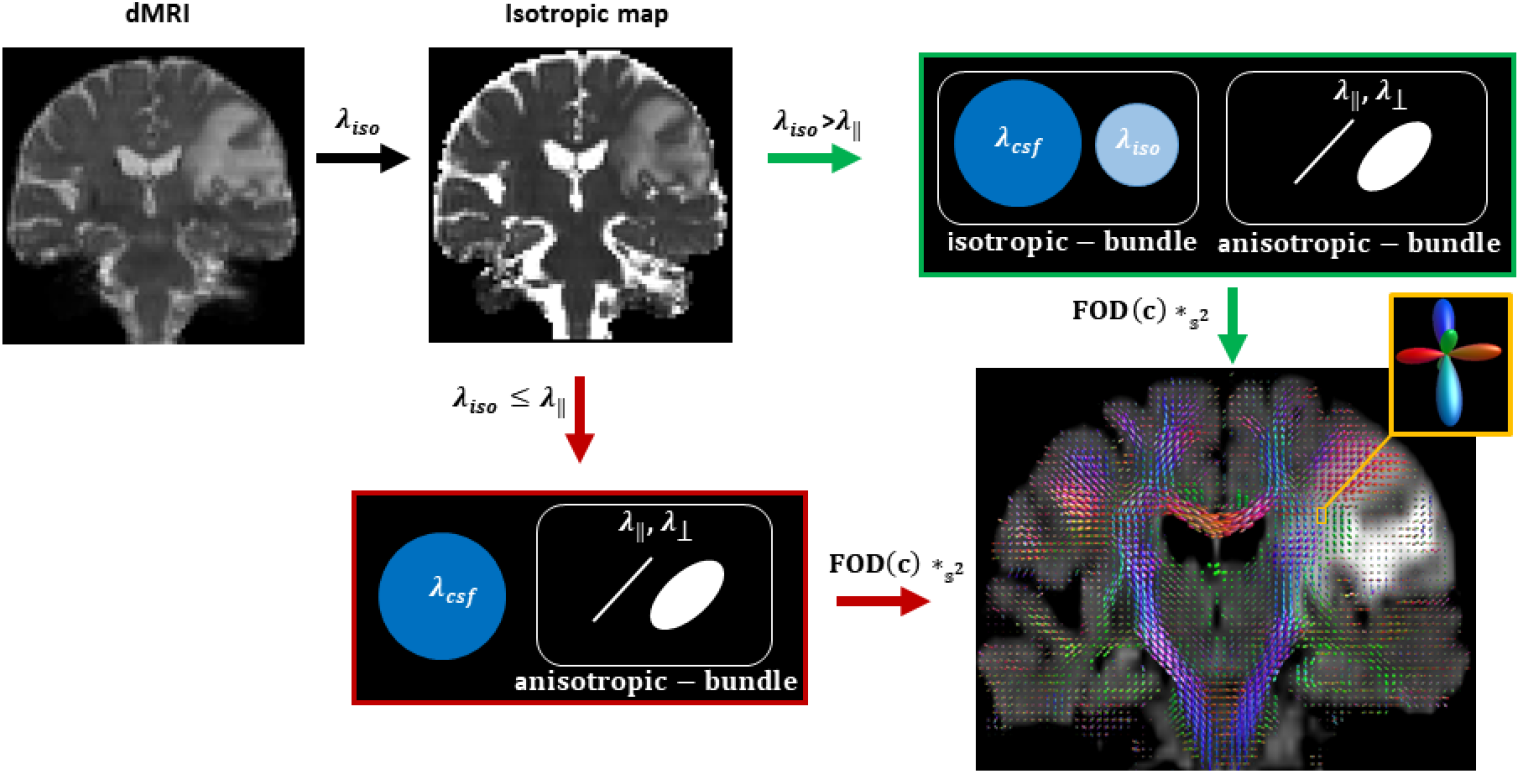
Schema of our proposed methodology. The green arrows show the fitting corresponding to WM voxels in the edematous region, and the red arrows demonstrate the fit in non-edematous tissue. The symbols were described in the text.

### Simulated data

To evaluate the accuracy of our proposed method using data with known ground truth for GM, WM, CSF and edema, we simulated a diffusion weighted imaging (DWI) signal using Phantomas [19] for a range of conditions and parameters, including volume fractions, SNR, fiber separation angles, and emulating single-shell and multi-shell data. Three protocols of single- and multi-shell data were simulated (Table.1). We demonstrated the complexity of our model containing GM, WM, CSF, and edema regions with crossings of two or three fibers which are surrounded by regions of isotropic diffusivity (Fig. 2). There were three regions of edema simulation, two with simulated fibers crossing at 90^°^ and 45^°^ and one with three fibers crossing at 90^°^ and 60^°^. Intrinsic diffusivity *λ*_∥_ of all simulated WM was set to 1.7*e* 9*m*^2^*/s*. Data were simulated with a range of isotropic volume fractions (20%, 40%, 60% and 80%) with *λ*_*iso*_ in a range of (1.7*e* −9*m*^2^*/s*, 2.95*e* − 9*m*^2^*/s*). Furthermore, Rician noise was added to the DWI signal, and our model was fit to the data either by *l*_*max*_ = 8, or *l*_*max*_ = 10, 12, 14, 16 using super-CSD. Each simulation was repeated 100 times. Inspired by the analysis in [20], we defined peak separation rate as the percentage of simulated voxels where the correct number of peak orientations was estimated. The algorithm identified additional peaks if the amplitudes are at least 25% of the highest peak. We further calculated the error between the ground truth and estimated fiber crossing angles. Further experiments with simulated data with b-values up to 3000 *s*/*mm*^2^ and number of gradient directions up to 256 are described in Supplementary Material.

**Table 1.** The list of evaluation protocols. The b-value is in units of *s*/*mm*^2^ and the number of simulated directions is shown in parentheses

| Protocol | b-value (DWI direction) |
| --- | --- |
| P1 | b=800(30) |
| P2 | b=2000(64) |
| P3 | b=300(15), b=800(30), b=2000(64) |

**Fig 2.**
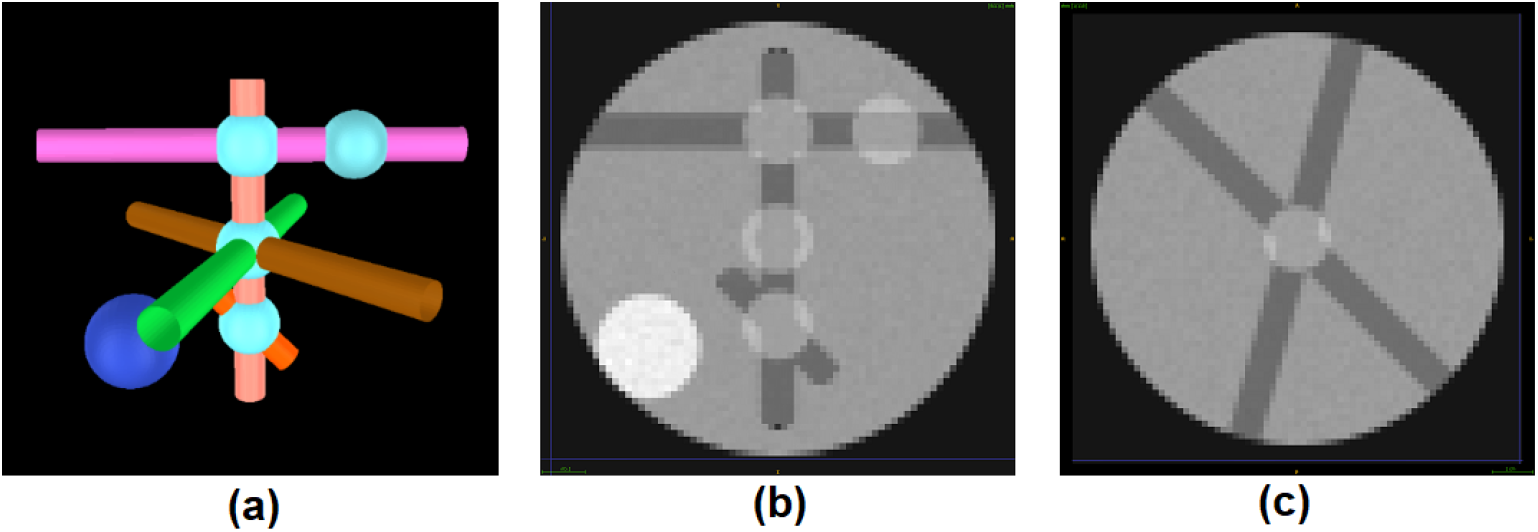
Simulated data with Phantomas. (a) The 3D visualization of Phantomas simulation, with GM (black), WM (tubes), edema regions surrounding crossings of two and three fibers with low isotropic diffusivity (cyan), and CSF (dark blue). (b) and (c) depict the coronal and axial view of a simulated diffusion weighted image, respectively.

### *In vivo* data acquisition, preprocessing and experiments

Twenty-three participants were recruited between 2017 - 2020 from an ongoing brain tumors study with varying degrees of peritumoral edema, including 13 patients with a histological diagnosis of glioblastoma (WHO grade 4), 8 with anaplastic gliomas (WHO grade 3 atrocytomas and oligodendrogliomas), and 2 with brain metastasis. Authors had access to information that could identify individual participants during or after data collection. All participants underwent a multi-shell diffusion acquisition with b-values of 300 (15 directions), 800 (30 directions), and 2000 (64 directions) *s*/*mm*^2^ and 9 b=0 volumes, with TR/TE of 4300/75 ms, along with the pulse duration time *δ* = 0.01293 s and pulse separation time Δ = 0.03666 s, multi-band factor of 2 and *A* > *P* phase encoding direction, followed by an acquisition of 7 b=0 volumes with *P* > *A* phase encoding direction. Data were acquired on a Siemens 3T Magnetom Prisma Fit scanner with either a 32-channel or 64-channel head coil. The data were denoised [21], followed by correction for echo planar imaging distortions with TOPUP [22] and motion and eddy current correction with EDDY [23]. Additionally, each patient underwent a routine pre-surgical scan in the clinic on either a Siemens TrioTim, MAGNETOM Vida or MAGNETOM Avanto 3T scanner, yielding T1-weighted (pre- and post-gadolinium contrast), T2-weighted and FLAIR images. These structural images were processed with the pipeline described in [24] which included DeepMedic [25] segmentation of the tumor and peritumoral area. The T1 image and segmentation were finally registered to the multi-shell DTI acquisition with ANTs software [26].

To compare FODs in our model, we fitted COMPARI and MSMT-CSD in each patient. We calculated the maximum peak amplitude of the FOD in every voxel in the peritumoral region. To illustrate the degree of anisotropy in our model, we used the anisotropic index (AI), an alternative to FA for FODs, as described in [27]. In addition, we selected one patient with metastatic tumor and edema affecting the *centrum semiovale*, a region of the brain known to have WM fiber crossings, to visualize number of peaks and FODs in the peritumoral region, and to visually compare results of COMPARI in edema to the contralateral WM. In the same patient, we fitted COMPARI and MSMT-CSD [10] as well as NODDI [28], and the multi-shell two-compartment model for free-water elimination from Hoy et al [29]. We visualized orientations of FOD peaks or tensor orientations between these models in the peritumoral region. Finally, we performed whole brain probabilistic tractography with both models, using the IFOD2 algorithm [30] and] and the SIFT model to determine seed points dynamically [31] in our selected patient. The ‘cutoff’ parameter for COMPARI was 0.2, while it was the default 0.1 for the MSMT-CSD model. It is noteworthy that the term ‘cutoff’ refers to the amplitude threshold of the FOD, and increasing the ‘cutoff’ value leads to more terminated streamlines. Four tracts arcuate fasciculus (AF), corticospinal tract (CST), corpus callosum (CC), and superior longitudinal fasciculus III (SLF III) of the peritumoral region were manually reconstructed by a neurosurgeon experienced in structural connectivity using Trackvis software (Department of Radiology, Massachusetts General Hospital, Boston, MA, USA).

### Implementation

We implemented COMPARI using the Dmipy [13] software package. COMPARI was fitted by employing a brute force optimization technique to sample all possible parameter values within the defined optimization bounds, followed by the application of constrained optimization algorithm L-BFGS-B [32] to obtain the locally optimal parameter estimates. Then, the CSD estimation was employed based on the approach proposed by [10]. The *λ*_∥_ initiated using response function described in [10]. In addition, we utilized Dipy [33], ITK-snap [34], MRview [35], and Trackvis [36] to facilitate visualization. We fitted COMPARI in parallel in each axial slice on a cluster with 40 cores 1.2 GHz CPU with 128 GB of RAM, and computation time was approximately 5 hours per participant.

## Results

We assessed our algorithm in a simulation setup with different parameters. Then, the tuned parameters were utilized in in vivo experiments.

### Simulations

We first assessed fiber crossings at angles of 90^°^, 60^°^, and 45^°^ in the context of simulated peritumoral edema. The peak separation rate and error of FOD separation angles were plotted for P1 and P2 (single-shell) protocols, with SNR=30 and *l*_*max*_= 8 using CSD method and isotropic volume fractions vf20%, vf40%, vf60%, and vf80% (Fig.3).

**Fig 3.**
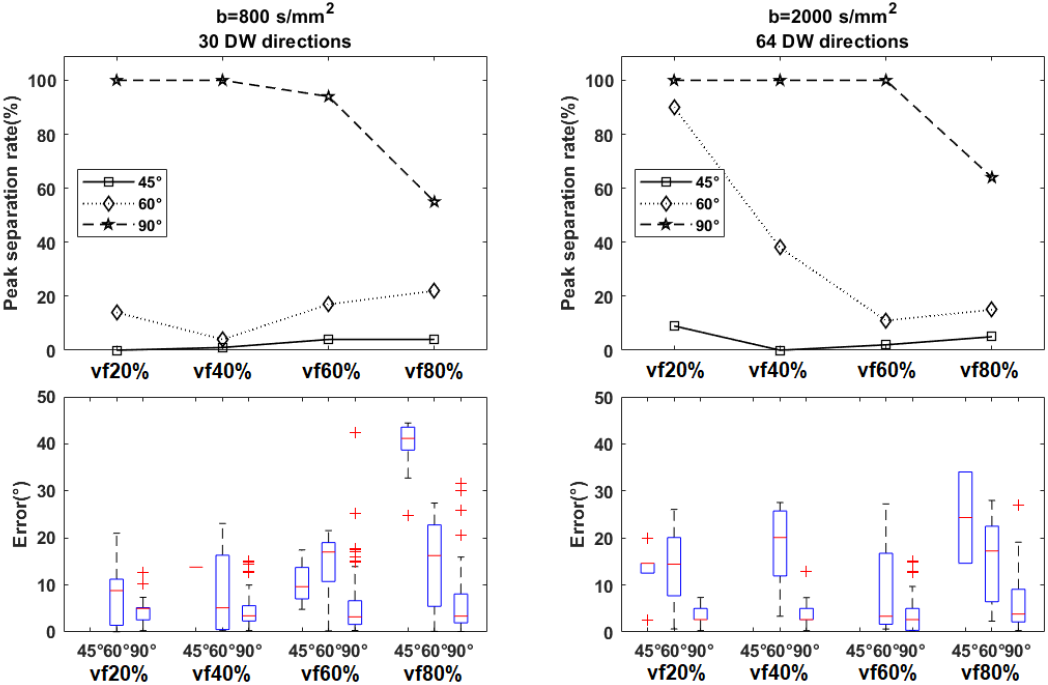
Simulation results of the effect of b-value and number of gradient directions on the ability to resolve crossing fibers. (first column: b=800 *s*/*mm*^2^(P1), second column: b=2000 *s*/*mm*^2^(P2). For all experiments, SNR=30 and *l*_*max*_ = 8. Peak separation rate for 60^°^ and 90^°^ crossings were the best in the P2 protocol with b=2000 *s*/*mm*^2^ Top: the peak separation rate of recovering peaks for ground truth crossing angles of (45^°^,60^°^,90^°^) with various isotropic volume fractions denoting different extent of peritumoral edema. Bottom: the distribution of errors in the crossing fiber angle are shown as boxplots.

We assessed the peak separation rate and error in crossing fiber angle with *l*_*max*_ = 8, 10, 12, 14 and 16 at SNR=30 in P3 protocol (Fig.4). We observed a significant improvement in resolving smaller separation angles with *l*_*max*_ > 8 compared to *l*_*max*_ = 8. In addition, the peak separation rate and error in crossing angle were improved for 60^°^crossings; however, there was no remarkable improvement for 90^°^ crossings, nor for *l*_*max*_ > 12. As a result, the super-CSD method with *l*_*max*_ = 12 was selected for subsequent experiments in human data. Further experiments with simulated data showed increased peak separation rate and reduced error with higher b-values and number of gradient directions (Supplementary Material).

**Fig 4.**
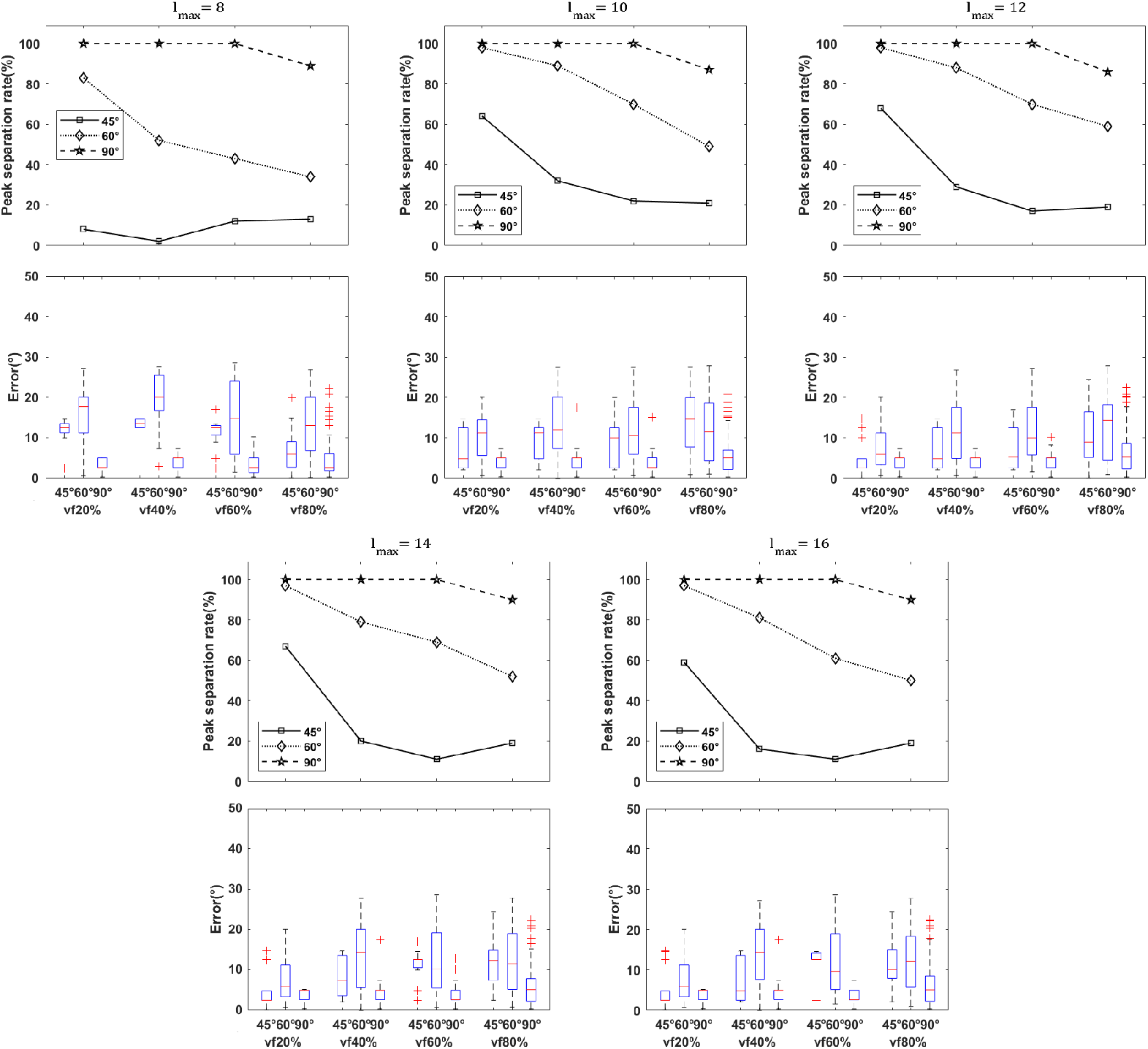
Simulation results of the effect of super-CSD fitting and edema on the ability to resolve crossing fibers in simulated multi-shell protocol (P3). For all experiments, the SNR was 30. Among super-CSD approaches, results are similar, with *l*_*max*_ = 12 showing highest peak separation rate for the 60^°^ crossing. Top: the peak separation rate of recovering peaks for ground truth crossing angles of (45^°^,60^°^,90^°^) with various isotropic volume fractions denoting different extent of peritumoral edema, for super-CSD fits with *l*_*max*_ = 8, 10, 12, 14 and 16. Bottom: the distribution of errors in the crossing fiber angle are shown as boxplots.

Furthermore, we separately assessed P3 protocol (multi-shell), with SNR= 20, 30, 40 and *l*_*max*_ = 12 (Fig. 5). As we expected, increasing Rician noise reduced the peak separation rate for crossings at all levels of VF. In addition, results in P3 protocol (Fig.5) and in the P2 protocol (Fig. 3) at SNR of 30 showed that the peak separation rates for 90^°^ and 60^°^ crossings were better in the P3 protocol compared to the P2 protocol at all levels of isotropic volume fraction, suggesting that COMPARI is more robust to extracellular water when using multi-shell data.

**Fig 5.**
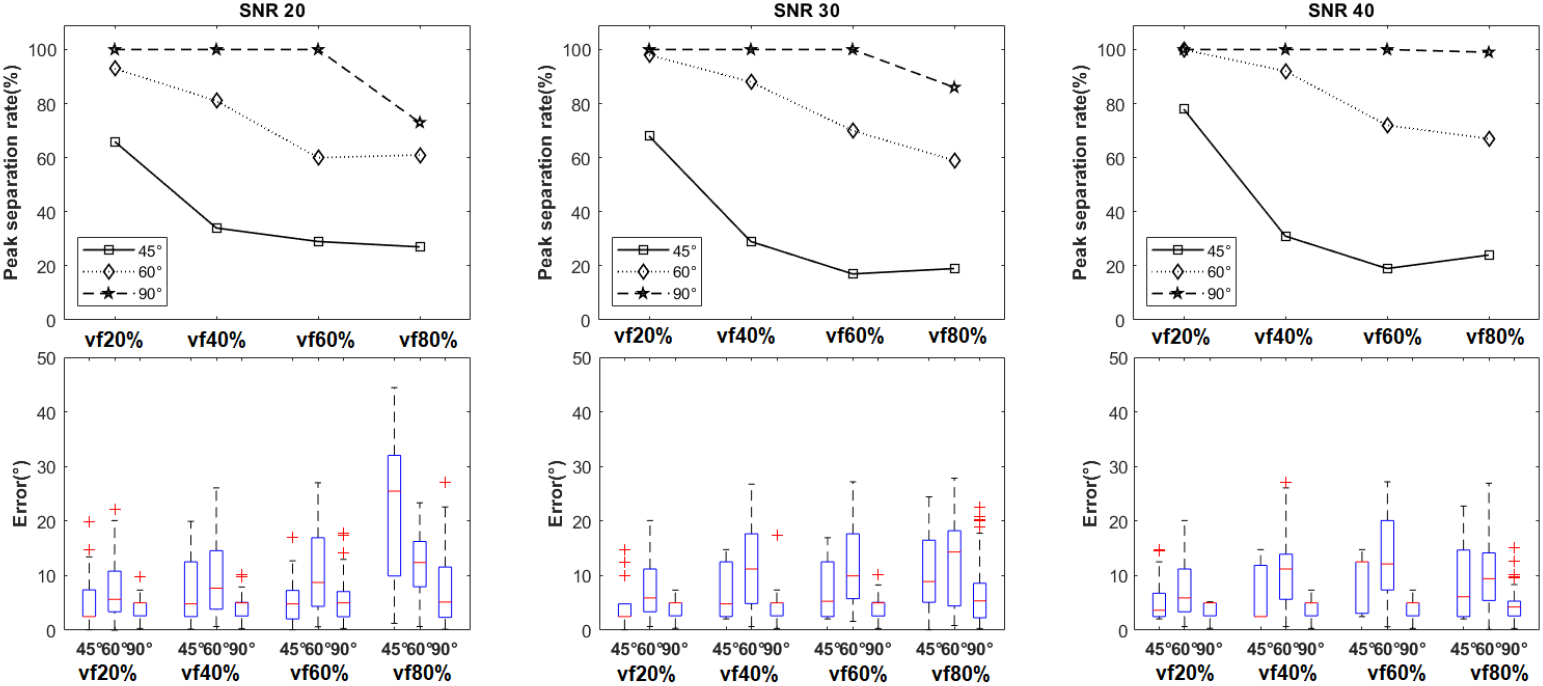
Simulation results of the effect of SNR and edema on the ability to resolve crossing fibers. For all experiments, the multi-shell protocol (P3) was used and *l*_*max*_ = 12. Higher SNR is associated with improved peak separation rate and error. Top: the peak separation rate of recovering peaks for ground truth crossing angles of (45^°^,60^°^,90^°^) with various isotropic volume fractions denoting different extent of peritumoral edema at three levels of Rician noise (SNR= 20, 30, 40). Bottom: the distribution of errors in the crossing fiber angle are shown as boxplots.

### In vivo experiments

#### Comparison of FOD amplitude and AI

Maximum amplitude of FODs and AI distribution within the peritumoral region obtained with COMPARI were compared to those extracted with the MSMT-CSD models (Fig.6). Median amplitudes as well as median AI were higher in COMPARI than in MSMT-CSD in all patients of the clinical dataset, with average FOD amplitude of 0.10 in MSMT-CSD compared to 0.54 in COMPARI.

**Fig 6.**
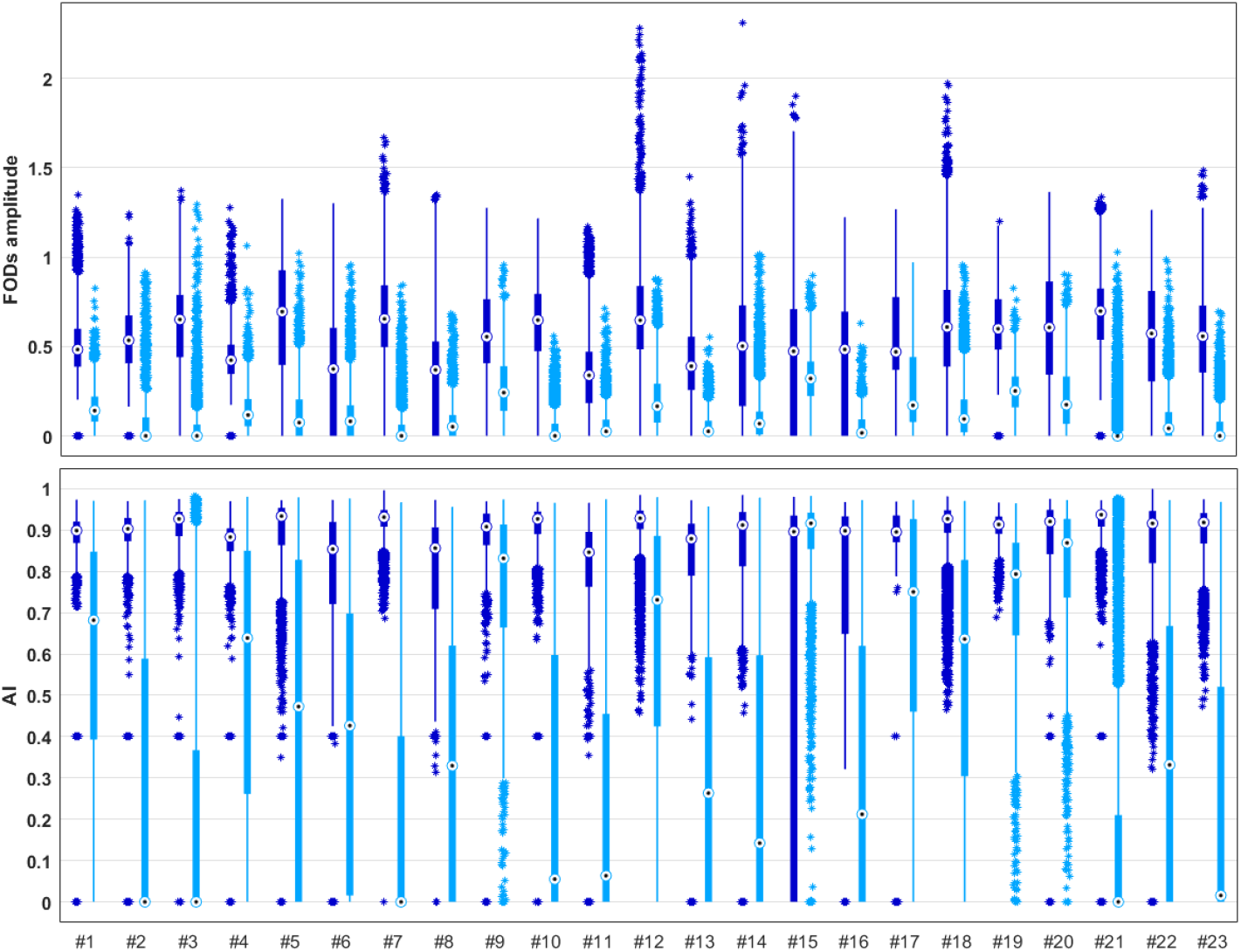
Strip plots of FOD amplitudes and AI in the edema region for all patients in our dataset. 23 patients are shown along the x-axes. Top: The distribution of COMPARI FOD peak amplitude is shown in dark blue and that of MSMT-CSD depicted in light blue. Bottom: The distribution of AI is shown in dark blue for COMPARI and that of MSMT-CSD depicted in light blue. Black dots inside circles ⊙ represent the median of each distribution.

#### FOD Visualization of COMPARI

COMPARI was applied in the peritumoral region as well as in the contralateral healthy homologous WM in a selected patient with brain metastasis (Fig.7). The number of FOD peaks and the amplitude and orientation were similar between the peritumoral and contralateral regions. The selected voxels in the peritumoral region (centrum semiovale) and contralateral side (healthy WM) demonstrated similarity of orientation and crossing FODs.

**Fig 7.**
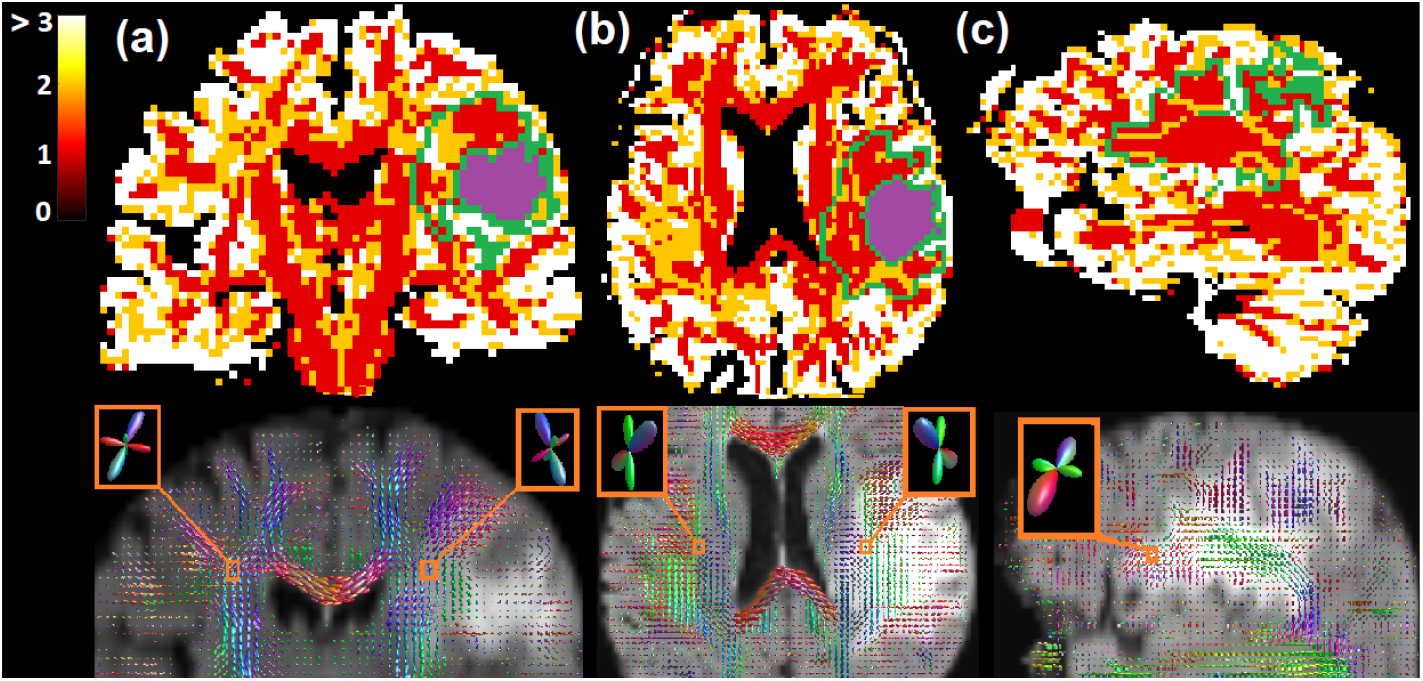
Results of COMPARI fitting in the peritumoral and contralateral *centrum semiovale*. Top: The map of the number of peaks representing crossing fiber populations per voxel. Voxels with more than 3 peaks are depicted in white. The edema region is outlined in green, and the tumor is in purple. Bottom: FODs visualized on a background of FLAIR in the (a) coronal plane, (b) axial plane, and (c) sagittal plane.

#### Visualization of multi-shell models

We visually compared orientation maps of MSMT-CSD with COMPARI in edema region. Additionally, we applied the NODDI [28] and HOY et al [29] approaches to our multi-shell data to compare the fitting of the edema region using multi-compartment and free-water approaches respectively(Fig.8). In NODDI (Fig.8a) and in the Hoy model (Fig.8b), no fiber crossings were captured. While MSMT-CSD (Fig.8c) FODs showed some crossings in the *centrum semiovale*, the difference in amplitude between FODs inside and outside the edema region was clearly visible. Finally, COMPARI (Fig.8d) reconstructed crossings in the edematous *centrum semiovale* with FOD amplitudes similar to those outside the edema.

**Fig 8.**
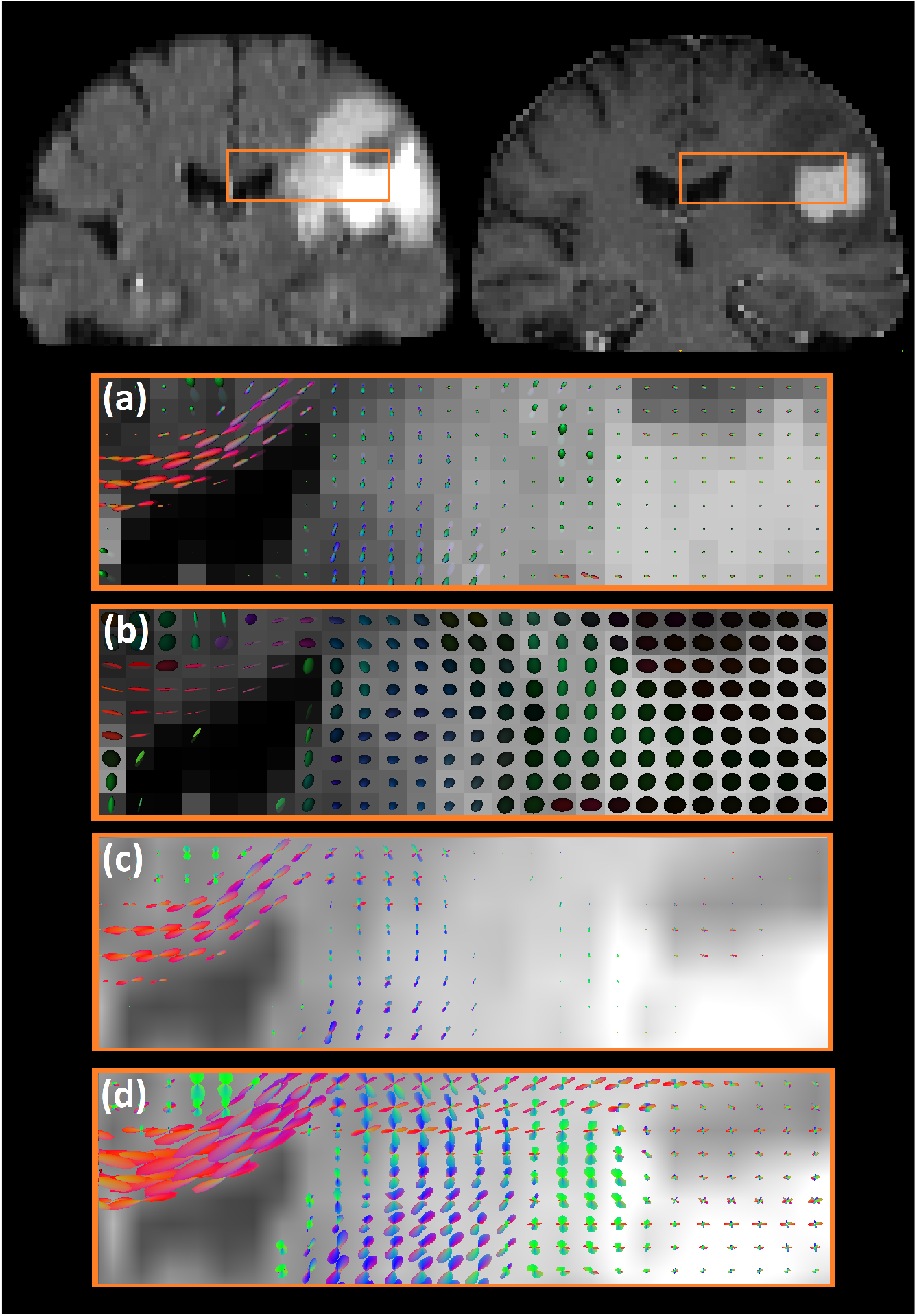
The comparison of orientation maps between multi-shell models. The top left and top right images are coronal slices of FLAIR and T1 contrast-enhanced images, respectively. The selected area covers the corpus callosum, *centrum semiovale*, peritumoral edema, and tumor regions. Orientation maps are shown on a background of FLAIR. (a) NODDI orientation dispersion maps; (b) tensors fit with Hoy et al free-water elimination model; (c) and (d) depict FODs of MSMT-CSD and COMPARI, respectively.

#### Tractography comparison between FOD models

COMPARI-based tractography was compared to that of MSMT-CSD (Fig.9). Streamlines of the CC partially crossed the CST using the MSMT-CSD (Fig.9a). AF and SLF III fibers located in the edema could not be visualized. In contrast, the crossing of the CST and CC fibers could be visualized with COMPARI, with a better qualitative definition of the CST (Fig.9b). Additionally, the AF and SLF III were visualized crossing each other in the peritumoral region with COMPARI.

**Fig 9.**
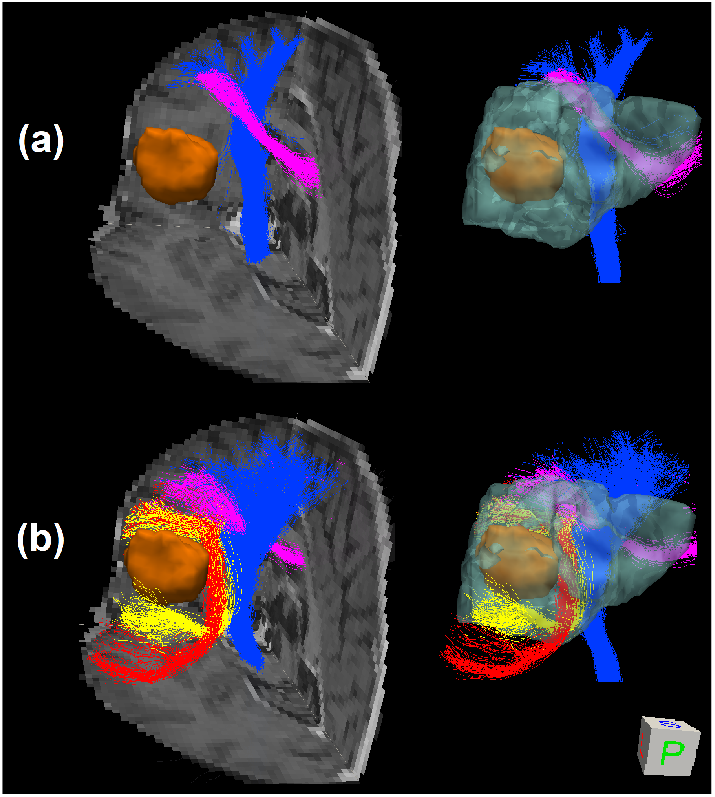
3D visualization of probabilistic tractography from the posterior vantage point. using (a) MSMT-CSD and (b) COMPARI, showing part of the CC (pink), CST (blue), AF (red) and SLF III (yellow). Tracts are shown overlaid on three slices of the T1 image on the left, and without a background on the right with the edema depicted in translucent cyan. The tumor is shown in orange in both views.

## Discussion

In this study, we introduced a multi-compartment modeling approach (COMPARI) to resolve crossing fibers in the peritumoral edema of brain tumor patients. Simulated and in vivo experiments show that crossing fibers were enhanced in edematous regions.

In simulation results (Fig.3-5), we tested the effects of b-value, number of DW directions, SNR, volume fraction and spherical harmonic order parameter (*l*_*max*_) on the ability of our model to detect crossing fibers. As shown in Fig. 3, our model could resolve angles of 60^°^ to 90^°^ with increasing success as b-value and number of DW directions increased. Additionally, results of the simulated multi-shell protocol (P3) outperformed those of single-shell protocols (P1, P2), suggesting that multi-shell data, including a low b-value to estimate the isotropic diffusivity and a high b-value with a large number of DW directions to resolve angular separation, is a prerequisite for our model. In results on simulated multi-shell data using CSD fitting (Fig.5), as expected, increased SNR was associated with higher peak separation rate in resolving crossing fibers, although this effect was small. We could not resolve fibers crossing at 45^°^, in line with [37]. However, with the super-CSD approach (Fig.4), we could resolve crossings as low as 45^°^ when volume fraction was as low as 20%. In higher volume fractions, as expected, the ability to resolve crossing fibers decreased, but our model could resolve crossings of 90^°^ with nearly 100% peak separation rate and a median of approximately 5^°^ of error. It could also resolve 60^°^ crossings, albeit with a lower peak separation rate.

For all values of *l*_*max*_ tested, the super-CSD fit (*l*_*max*_ > 8) outperformed the CSD fit, with not much difference between values of *l*_*max*_. Additional simulated experiments (S1 Table) showed that increasing b-values and number of gradient directions improved peak separation rate (Supplementary Material, S1 Fig-S20 Fig), although b-values of 3000 *s*/*mm*^2^ and 256 directions may not be feasible with clinical scanners.

In human data, there is no ground truth to assess the quality of COMPARI fit. Metastatic brain tumors are known to have the lowest level of tumor infiltration in the peritumoral area [38] with a high degree of edema. Accordingly, we compared fiber crossings in the edematous peritumoral area near the *centrum semiovale* of one patient with brain metastasis to the contralateral homologous area. We found that COMPARI model was able to resolve fiber crossings similarly in both hemispheres (Fig. 7). Upon comparison with other multi-shell models (Fig.9), we found that crossing fibers in edema with non-negligible amplitude were only identified using COMPARI. Since the amplitude cutoff is a parameter essential for tractography, corresponding results of COMPARI were considerably improved over MSMT-CSD (Fig. 9). To ensure that this increase in amplitude is a generalizable result, we repeated this in 22 other patients, and found that COMPARI FODs consistently had higher amplitude and higher AI than those of MSMT-CSD (Fig. 6).

The diffusion kurtosis effect [39] is known to cause inaccurate estimation of diffusivity using Gaussian diffusion models at b-values higher than b=1500 *s*/*mm*^2^. As a result, our model performed poorly on a human data set with b=3000 *s*/*mm*^2^ and 128 directions (S21 Fig), since it relies on an estimate of isotropic diffusivity. This effect was not seen in simulated data due to a limitation of the simulation model. Thus, from our experiments on simulated and human data, we concluded that best results with COMPARI would be achieved using a multi-shell dataset with a low b-value shell to capture isotropic diffusivity, and one or more high-b shells to resolve fiber directions, fit with *l*_*max*_ = 12 with the super-CSD approach, if possible.

Results of probabilistic tractography on a patient with glioblastoma showed that COMPARI was able to replicate the results of MSMT-CSD of the CC intersecting the CST, with a better qualitative definition of the CST, while further elucidating two crossing fiber bundles in the edema region (AF and SLF III) which were completely missed when performing tractography with the MSMT-CSD model. This result is probably a combination of higher FOD peak amplitude with COMPARI, as well as a better ability to separate edema from the underlying WM.

In addition to the comparison of our proposed model, COMPARI, and MSMT-CSD, we also conducted further experiments and comparisons, which are presented in the Supplementary Material. These experiments include a comparison of the normalized FODs obtained from COMPARI and MSMT-CSD, as shown in S22 Fig, which showed that the low amplitudes of FODs fit from the MSMT-CSD model cannot be compensated for with simple normalization. We also compared the deterministic tractography results obtained from FERNET and COMPARI, where only the latter was able to distinguish the AF from the SLF III (S23 Fig). Additionally, we compared a simpler model of COMPARI described in Supplementary Material to the full COMPARI model to demonstrate the necessity of the full model (S24 Fig).

There are several limitations of this study. First, our model is based on several constraints, including the tortuosity constraint and equality of the *λ*_∥_ value for stick and zeppelin compartments, that were made with the goal of modeling crossing fibers in clinically feasible data. Therefore, derived values such as volume fractions should not be ascribed an explicit biological meaning, although they may be useful as clinical markers. A second potential limitation of our method is that it could fail in the grey matter (GM) if isotropic diffusivity is higher than *λ*_∥_. In this case, FODs fitted to the GM had an excessive number of peaks (Fig. 7), which could result in false positives in tractography connecting GM regions. As such, care should be taken to fit COMPARI only to the WM, or to restrict tractography to the WM and peritumoral area, such as with anatomically constrained tractography [40, 41]. Third, while simulated results suggest that best results with COMPARI will be achieved with multi-shell datasets, COMPARI has not yet been tested in *in vivo* multi-shell datasets with a variety of b-values and direction sets.

In clinical practice, tractography could be useful for clinicians treating patients with brain tumors, especially in the context of surgical planning. Indeed, tumor resection is the first treatment for most tumor types [42], and the quality of the resection (i.e. amount of tumor removed) is the main factor under consideration [43], especially for gliomas where maximal safe resection is the standard of care [44]. Particular attention must nevertheless be paid to structural connectivity (i.e. WM tracts), in order to avoid new neurological deficits and/or neurological impairments in patients, and thus preserve their quality of life [45]. The identification - and consequently the preservation - of WM tracts during brain surgery is therefore of major importance, for both oncological and functional reasons.

The presence of edema has been a confounding variable [46], and the application of the COMPARI protocol to surgical tumor cases should improve the accuracy of tractography in peritumoral areas, to help achieve onco-functional balance [47], neurological preservation, and maximal safe resection.

## Conclusion

Our study we present an improved crossing fiber model (COMPARI) for clinically feasible dMRI data, which surpasses MSMT-CSD in its ability to model WM in the presence of excessive free water. Through the removal of isotropic compartments, we are able to significantly improve crossing fiber modeling in edema, enabling tractography through complex WM structures in the peritumoral area. Through the application on brain tumor data, we have shown its clinical applicability in improving surgical planning.

## Data availability

The datasets generated during the current study are not publicly available due to the IRB requirements of the Hospital of University of Pennsylvania. The datasets can be made available on request to the corresponding author, after required data transfer and IRB paperwork is completed. The code is available here: https://github.com/egolkar/COMPARI.

## Ethical Standards

The experimental protocol was reviewed and approved by the Institutional Review Board of the University of Pennsylvania. All participants were recruited from the University of Pennsylvania and provided written informed consent.

## Funding

The current work was supported in part by NIH grant RO1 NS096606 (PI, Drs Verma and Brem) and by a research grant from Synaptive Medical Inc.

## Disclosures

The authors have no personal, financial, or institutional interest in any of the drugs, materials, or devices described in this article.

## Supplementary Material

**S1 Table. Further simulated data protocols**.

**S1 Fig. Simulation results of data with b=800** *s*/*mm*^2^ **and 64 directions, as**

*l*_*max*_ **is increased from 8 to 10 and 12**.

**S2 Fig. Simulation results of data with b=2000** *s*/*mm*^2^ **and 64 directions, as** *l*_*max*_ **is increased from 8 to 10 and 12**.

**S3 Fig. Simulation results of data with b=3000** *s*/*mm*^2^ **and 64 directions, as** *l*_*max*_ **is increased from 8 to 10 and 12**.

**S4 Fig. Simulation results of data with b=800-2000** *s*/*mm*^2^ **and 30-64 directions, as** *l*_*max*_ **is increased from 8 to 10 and 12**.

**S5 Fig. Simulation results of data with b=800-2000** *s*/*mm*^2^ **and 30-64 directions, as** *l*_*max*_ **is increased from 8 to 10 and 12**.

**S6 Fig. Simulation results of data with b=800** *s*/*mm*^2^ **and 91 directions, as**

*l*_*max*_ **is increased from 12 to 14 and 16**.

**S7 Fig. Simulation results of data with b=2000** *s*/*mm*^2^ **and 91 directions, as** *l*_*max*_ **is increased from 12 to 14 and 16**.

**S8 Fig. Simulation results of data with b=3000** *s*/*mm*^2^ **and 91 directions, as** *l*_*max*_ **is increased from 12 to 14 and 16**.

**S9 Fig. Simulation results of data with b=800-2000** *s*/*mm*^2^ **and 30-91 directions, as** *l*_*max*_ **is increased from 12 to 14 and 16**.

**S10 Fig. Simulation results of data with b=800-3000** *s*/*mm*^2^ **and 30-91 directions, as** *l*_*max*_ **is increased from 12 to 14 and 16**.

**S11 Fig. Simulation results of data with b=800** *s*/*mm*^2^ **and 128 directions, as** *l*_*max*_ **is increased from 12 to 14 and 16**.

**S12 Fig. Simulation results of data with b=2000** *s*/*mm*^2^ **and 128 directions, as** *l*_*max*_ **is increased from 12 to 14 and 16**.

**S13 Fig. Simulation results of data with b=3000** *s*/*mm*^2^ **and 128 directions, as** *l*_*max*_ **is increased from 12 to 14 and 16**.

**S14 Fig. Simulation results of data with b=800-2000** *s*/*mm*^2^ **and 30-128 directions, as** *l*_*max*_ **is increased from 12 to 14 and 16**.

**S15 Fig. Simulation results of data with b=800-3000** *s*/*mm*^2^ **and 30-128 directions, as** *l*_*max*_ **is increased from 12 to 14 and 16**.

**S16 Fig. Simulation results of data with b=800** *s*/*mm*^2^ **and 256 directions, as** *l*_*max*_ **is increased from 12 to 14 and 16**.

**S17 Fig. Simulation results of data with b=2000** *s*/*mm*^2^ **and 256 directions, as** *l*_*max*_ **is increased from 12 to 14 and 16**.

**S18 Fig. Simulation results of data with b=300** *s*/*mm*^2^ **and 256 directions, as** *l*_*max*_ **is increased from 12 to 14 and 16**.

**S19 Fig. Simulation results of data with b=800-2000** *s*/*mm*^2^ **and 30-256 directions, as** *l*_*max*_ **is increased from 12 to 14 and 16**.

**S20 Fig. Simulation results of data with b=800-3000** *s*/*mm*^2^ **and 30-256 directions, as** *l*_*max*_ **is increased from 12 to 14 and 16**.

**S21 Fig. Patients results of data with b=3000** *s*/*mm*^2^ **and b=5000** *s*/*mm*^2^

**S22 Fig. Displays a comparison of the fiber orientation distribution (FOD) maps between the normalized FOD using MSMT-CSD and the COMPARI method**.

**S23 Fig. Visualization of FOD orientation using a) the simpler model and b) the COMPARI method**.

**S24 Fig. Deterministic tractography visualization from a posterior viewpoint**.

## Notes

### Competing Interest Statement

The authors have declared no competing interest.

